# Galectin-3 and *N*-acetylglucosamine promote myogenesis and improve skeletal muscle function in the *mdx* model of Duchenne muscular dystrophy

**DOI:** 10.1101/203653

**Authors:** Ann Rancourt, Sébastien Dufresne, Guillaume St-Pierre, Julie-Christine Lévesque, Haruka Nakamura, Yodai Kikuchi, Masahiko S. Satoh, Jérôme Frenette, Sachiko Sato

## Abstract

The muscle membrane, sarcolemma, must be firmly attached to the basal lamina. The failure of proper attachment results in muscle injury, which is the underlying cause of Duchenne muscular dystrophy (DMD), where mutations in the dystrophin gene disrupts the firm adhesion. In DMD patients, even moderate contraction causes damage, leading to progressive muscle degeneration. The damaged muscles are repaired through myogenesis. Consequently, myogenesis is highly active in DMD patients, and the repeated activation of myogenesis leads to the exhaustion of the myogenic stem cells. Therefore, approaches to reducing the risk of the exhaustion are to develop a treatment that strengthens the interaction between the sarcolemma and the basal lamina, and increases the efficiency of myogenesis. Galectin-3 is an oligosaccharide-binding protein and known to be involved in cell-cell interactions and cell-matrix interactions. Galectin-3 is expressed in myoblasts and skeletal muscle while its function in muscle remains elusive. In this study, we found evidence that galectin-3 and the monosaccharide *N*-acetylglucosamine, which increases the ligands (oligosaccharides) of galectin-3, promotes myogenesis *in vitro*. Moreover, in the *mdx* mouse model of DMD, treatment with *N*-acetylglucosamine increased the muscle force production. Our results demonstrate that treatment with N-acetylglucosamine can mitigate the burden of DMD.

## Introduction

Galectin-3 is a member of the galectin family of soluble non-glycosylated proteins. Galectins share primary sequence homology in their carbohydrate-recognition domains (CRD), which have affinity for oligosaccharides that contain b-galactoside, where a galactose residue (Gal) is linked to a saccharide (typically *N*-acetylglucosamine, GlcNAc) via a β-linkage (1–3). Galectin-3 contains two domains: a C-terminal CRD and an N-terminal collagen-like domain consisting of a repeating peptide sequence rich in proline and glycine residues (4). Galectin-3 is expressed in many tissues, including muscle and one subset of pro-regenerative macrophages (M2), which are required to initiate muscle repair including adult myogenesis (5–8). Galectin-3 binds to a specific type of oligosaccharide attached to some membrane and extracellular glycoproteins, including cell adhesion molecules, integrins, some growth factor receptors, such as insulin-like growth factor receptor, and the major component of the basal lamina, laminin (9–12). Upon binding to the ligands, galectin-3 molecules oligomerize themselves through the N-terminal collagen-like domains. Consequently, the self-oligomerized galectin-3 becomes multivalent in oligosaccharide binding and directly cross-links its ligands on the cell surface or in the cell matrix (9, 13–15). When galectin-3 cross-links its ligand glycoproteins on the surface of a cell, it restricts the movements of those ligand glycoproteins (9). In contrast, when galectin-3 binds to its surface ligands on different entities, such as two different cells or cell and cell matrix, this ligand cross-linking results in cell-cell interactions or cell-matrix interactions (14, 16, 17). Through this cross-linking, galectin-3 exerts multifaceted activities, such as the modulation of signal transduction, membrane receptor dynamics, cell mobility, cell-cell adhesion, and cell-matrix interactions, all of which can be considered necessary steps in myogenesis. However, it remains unknown whether galectin-3, which is expressed in muscle, plays any role in myogenesis and in muscle function.

Myogenesis is a part of muscle regeneration processes, and is initiated in response to muscle fiber damage (18, 19). Growth factors are released from injured muscle cells and leukocytes, such as neutrophils and macrophages that are recruited to the site of injury. Those factors first trigger the differentiation of adult myogenic cells (satellite cells) into myoblasts. The myoblasts elongate, migrate, and make contact with neighboring myoblasts. As the cells adhere to one another, their membranes become intimately aligned and the myoblasts then fuse together to form multinucleated syncytial myotubes. The mechanism by which mammalian myoblasts undergo fusion remains elusive (20). Nascent myotubes then further fuse with myoblasts to form myofibers. The regenerated myofibers are anchored to laminin in the basal lamina by dystrophin-associated glycoprotein complex (DGC) (21)

Duchenne muscular dystrophy (DMD) is an X-linked lethal muscular dystrophy (MD) affecting one in 3,500 males at birth. It is the commonest childhood form of MD, and accounts for 50% of all MD cases (22, 23). Patients usually die in their late 20s. DMD is caused by mutations in the dystrophin gene, which lead to the loss of the cytoskeletal protein dystrophin. Dystrophin connects the cytoskeletal actin filaments to the basal lamina by binding to F-actin through its N-terminus and to β-dystroglycan in DGC through its C-terminus (24). DGC binds to laminin in the basal lamina, forming firm attachment of muscle membrane to the basal lamina. This stable anchoring of the muscle fibers to the basal lamina prevents injury associated with normal repeated and unaccustomed eccentric contractions. However, in the muscles of DMD patients, whose expression of dystrophin is lacking, the myofibers are not properly fixed to the basal lamina thereby detach from the basal lamina during contraction, leading to sarcolemmal instability and damage. Fiber degeneration is then counterbalanced by myogenesis at the expense of adult myogenic cells. The constant degeneration of the muscle fibers in MD patients eventually overwhelms the intrinsic capacity for myogenesis. There are very few clinically available therapies for DMD that mitigate disease progression, such as corticosteroids, although some are now in clinical trials, including drugs that can bypass inherited mutations (23, 25). While gene and cell therapies are on the horizon, it is important to develop new therapeutic approaches for DMD. Two possible approaches for delaying the progression of MD are to strengthen the sarcolemmal attachments to protect them from contractions and to increase the efficiency of myogenesis. In this study, we first investigated whether the efficiency of myogenesis *in vitro* is increased by galectin-3 or the monosaccharide GlcNAc that increases the biosynthesis of galectin-3 ligands. Then we also studied whether treatment with GlcNAc mitigates DMD using a mouse model of DMD.

## Materials and Methods

### Reagents

Recombinant galectin-3 was purified from an extract of *Escherichia coli* that overexpressed galectin-3, with affinity chromatography using lactosyl-Sepharose (Sigma-Aldrich St. Louis, MO, USA), as described previously (16, 26, 27). The bound galectin-3 was released with phosphate-buffered saline (PBS (–)) containing 150 mM lactose, and lactose in the fraction containing galectin-3 was then removed with a HiPrep™ 26/10 Desalting Column (GE Healthcare, Pittsburgh, PA, USA). The eluate containing galectin-3 was passed through an ActiClean EtOx (Sterogen, Carisabad, CA, USA) column to ensure that the endotoxin level was < 1 pg/μg. The specific (oligosaccharide-binding) activity of galectin-3 was tested before each experiment by using it to induce hemagglutination. An anti-myosin heavy chain (MHC) antibody (MF20) was obtained from the Developmental Studies Hybridoma Bank at the University of Iowa (Iowa City, IA, USA). An anti-galectin-3 antibody (M3/38.1.2.8) was purified and biotinylated as previously published (16, 26, 27).

### Animals

Male *mdx* dystrophic mice (C57BL/10ScSn-*Dmd*^*mdx*^/J) were purchased from the Jackson Laboratory (Bar Harbor, ME, USA) and bred at our animal facility. Galectin-3-null mice (G3KO) were obtained from Core F of the National Institute of General Medical Sciences-supported Consortium of Functional Glycomics. Wild type (C57BL/6) and G3KO mice were bred and maintained at our facility. *N*-Acetylglucosamine (250 mg/kg bodyweight per day) was administered intraperitoneally to the *mdx* mice for 10 days on days 25-35 after birth. The control mice were injected daily with the same volume of PBS. At the end of the experiment, the mice were euthanized using intraperitoneal injection of 50mg/Kg pentobarbital. All animal breeding and experimental procedures were approved by the Laval University Research Centre Animal Care and Use Committee, the decisions of which were made in accordance with the Canadian Council on Animal Care Guidelines.

### Myogenesis

Mouse-derived myoblast cell line, C2C12 (American Type Tissue Culture Collection, Manassas, VA, USA) were cultured for propagation in Dulbecco’s modified Eagle’s medium (DMEM; high glucose, 4.5g/l) containing 2 mM glutamine, supplemented with antibiotics (100 units/ml penicillin and 100 μg/ml streptomycin) and 10% heat-inactivated fetal calf serum (HyClone-GE Healthcare), and is referred to as ‘growth medium’ (GM). For differentiation, C2C12 cells (2.3 × 10^4^ cells in 30 μl of GM) were first plated into a well of μ-Slide VI^0.4^ (ibidi, GmbH, Germany) and incubated at 37 °C for 30-40 min to allow the cells to adhere to the well. GM (120 μl) was then added and the cells were cultured under a constant stream of 95% air/5% CO_2_ gas for 18-20 h. To initiate differentiation, the medium was replaced with DMEM containing 1% heat-inactivated horse serum (Sigma), antibiotics, insulin (10 μg/ml), transferrin (5.5 μg/ml), and sodium selenite (5 ng/ml), which is referred to as ‘differentiation medium’ (DM). The medium was changed every day after the initiation of differentiation. Under these conditions, elongated myotubes typically began to appear after 48 h, and some became thick mature myotubes after 72 h. The progression of myogenesis was estimated by either counting the number of nuclei or measuring the areas of MHC-positive and multinuclear myotubes using an inhouse software. The cells were treated with galectin-3 or GlcNAc, and the medium was changed every day after the initiation of differentiation.

### Immunofluorescent staining

The cells were fixed for 15 min in 3.7% paraformaldehyde in PBS (pH 7.4). For MHC staining, the fixed cells were treated with PBS containing 0.25% Triton X-100 for 5 min at room temperature, and then incubated with an anti-MHC antibody (2 μg/ml) for 1 h. After the cells were washed three times with PBS, they were incubated with an Alexa-Fluor-488 or 546-conjugated anti-mouse IgG antibody (ThermoFisher Scientific, Waltham, MA, USA) for 1 h. For galectin-3 staining, the cells were incubated with a biotinylated anti-galectin-3 antibody (2.5 μg/ml) for 1 h, and then with streptavidin-Alexa 488 or 546 (ThermoFisher Scientific). The cell nuclei were counterstained with 4′,6-diamidino-2-phenylindole (DAPI, ThermoFisher). The images were captured with the Quorum WaveFx Spinning Disc Confocal System (Quorum Technologies Inc., ON, Canada) with a Leica microscope controlled by the Volocity v4.0 software. The Quorum WaveFX-X1 Spinning Disc Confocal System (Quorum Technologies Inc.) with an Olympus IX inverted microscope controlled by the MetaMorph software (Molecular Devices, Sunnyvale, CA, USA) was used to capture the images in multiple fields of view (FOVs), and the images were then stitched together with in-house software (28). For immunofluorescence staining of muscle tissue, a frozen 5 μm tissue section was exposed to cold acetone for 10 min, then exposed to 0.3% H_2_O_2_-PBS for 5 min, and washed in PBS. After incubation with blocking buffer containing 0.05% Tween, 0.2% gelatin, 3% BSA, and 2% horse serum in 100 mM Tris-HCl (pH 7.5) and 150 mM NaCl for 45 min, the sections were exposed to the biotinylated anti-galectin-3 antibody overnight at 4 °C. After the sections were washed with PBS, they were incubated with streptavidin-Alexa 488 for 1 h and the nuclei were stained with DAPI.

### Isometric contractile properties

Mice were intraperitoneally injected with 0.1 mg/kg bodyweight buprenorphine to palliate the poor analgesic effect of pentobarbital. After 15 min, the mice were anesthetized with 5 mg/kg bodyweight sodium pentobarbital. The soleus and extensor digitorum longus (EDL) muscles were dissected, incubated in buffered physiological salt solution (Krebs-Ringer), and attached to an electrode and a force sensor (305B-LR, Aurora Scientific, Aurora, ON, Canada) to assess their contractile properties, as described previously (29). The maximum tetanic tension (P0, g) values were obtained using the sensor controlled by dynamic muscle control and data acquisition software (Dynamic Muscle Data Analysis software version 6.1, Aurora Scientific). The maximum specific tetanic tension (sP0, N/cm^2^) was calculated by dividing the wet weight by the optimal muscle length, multiplied by the muscle density (1.06 g/cm^2^), multiplied by the fiber/muscle length ratio. After the contractile properties were measured, the tendons were removed and the muscles were weighed.

### Statistical analysis

Statistical significance was determined with one-way ANOVA. All statistical analyses were performed with Prism (GraphPad Software Inc., La Jolla, CA, USA), and differences were considered significant at P < 0.05.

## Results

### Galectin-3 is expressed in myoblasts and on the sarcolemmal membrane of skeletal muscle

C2C12 cells are murine myoblasts and can be committed to myogenesis by placing them from GM into low-serum DM. We first analyzed the expression of galectin-3 in differentiating murine myoblasts and myotubes. As shown in Fig. 1A, galectin-3 was expressed in both proliferating myoblasts and myotubes while its expression was significantly reduced 24 h after the initiation of differentiation. The release of galectin-3 to the medium was significantly increased 24 h after the initiation of differentiation and gradually declined thereafter (Fig. 1B). These results suggest that the synthesis and localization of galectin-3 is regulated during myogenesis. Immunostaining of galectin-3 also showed that galectin-3 is highly expressed in both differentiated myoblasts and thin nascent myotubes (Fig. 1C a-f, data not shown), whereas mature myotubes do not express high levels of galectin-3 (Fig. 1C g-q). In skeletal muscles, galectin-3 was mainly found on the sarcolemmal membranes (Fig. 1D).

**Fig. 1.**
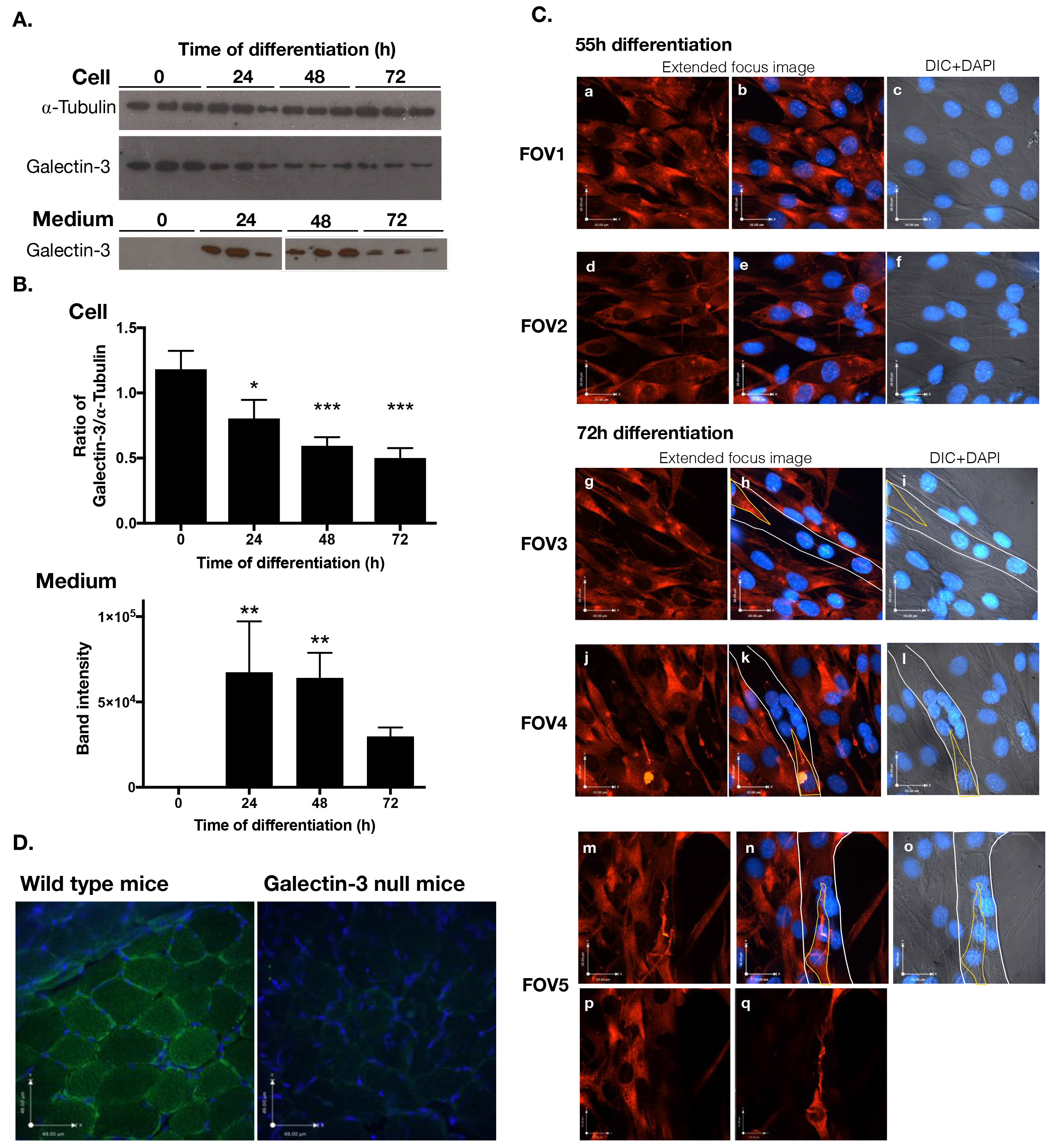
Expression of galectin-3 in differentiating myoblasts, myotubes, and muscles. A and B. Differentiation of murine myoblast C2C12 cells was induced by changing the medium from GM to DM. The levels of galectin-3 in the cells and medium were detected with western blotting using an anti-galectin-3 antibody. Results of densitometry analyses are shown in B. Significance compared to 0h of differentiation. * P < 0.05, ** P < 0.01, *** P < 0.001 C. Galectin-3 in myoblasts and myotubes 55h (FOV1 and 2) or 72h (FOV 3, 4 and 5) of differentiation was visualized with immunostaining using an anti-galectin-3 antibody (red). Extended-focus images of the fluorescence (red) and an optical plane image of stained nuclei (DAPI, blue) were merged (a, b, d, e, g, h, j, k, m, and n); a focal-plane DIC images and DAPI images are merged and shown (c, f, i, l, o). Myotubes were marked with thin white lines and myoblasts that attached onto the myotubes were marked with yellow lines in FOV3, 4 and 5. A z-optical image close to the bottom of the cover glass (p) and a z-optical image close to the top of a myotube of FOV5 (images m-o) show a lack of galectin-3 expression in the myotubes and the concentrations of galectin-3 on the extended myoblasts attached to myotubes.

### Galectin-3 promotes myoblast differentiation

Because the expression and release of galectin-3 are regulated during myogenesis, we next investigated whether galectin-3 increases the efficiency of myogenesis. First, the effect of an antagonist saccharide of galectin-3, lactose on the myogenesis was tested. The numbers of nuclei in myotubes were counted to estimate the progression of myogenesis. The myogenesis was significantly inhibited by the presence of lactose (Fig. 2A). Then we studied whether exogenously added galectin-3 further increases the formation of myotube. As little as 0.2 μM galectin-3 induced a 2.6-fold increase in the myogenesis (Fig. 2B). The myotubes that formed in the presence of galectin-3 contained more nuclei than those that formed in its absence (Fig. 2C). The stitched images of 7 × 5 FOVs showed that in the presence of galectin-3, the myotubes were longer (some exceeded 1 mm) than those formed in its absence (Fig. 2D and E).

**Fig. 2.**
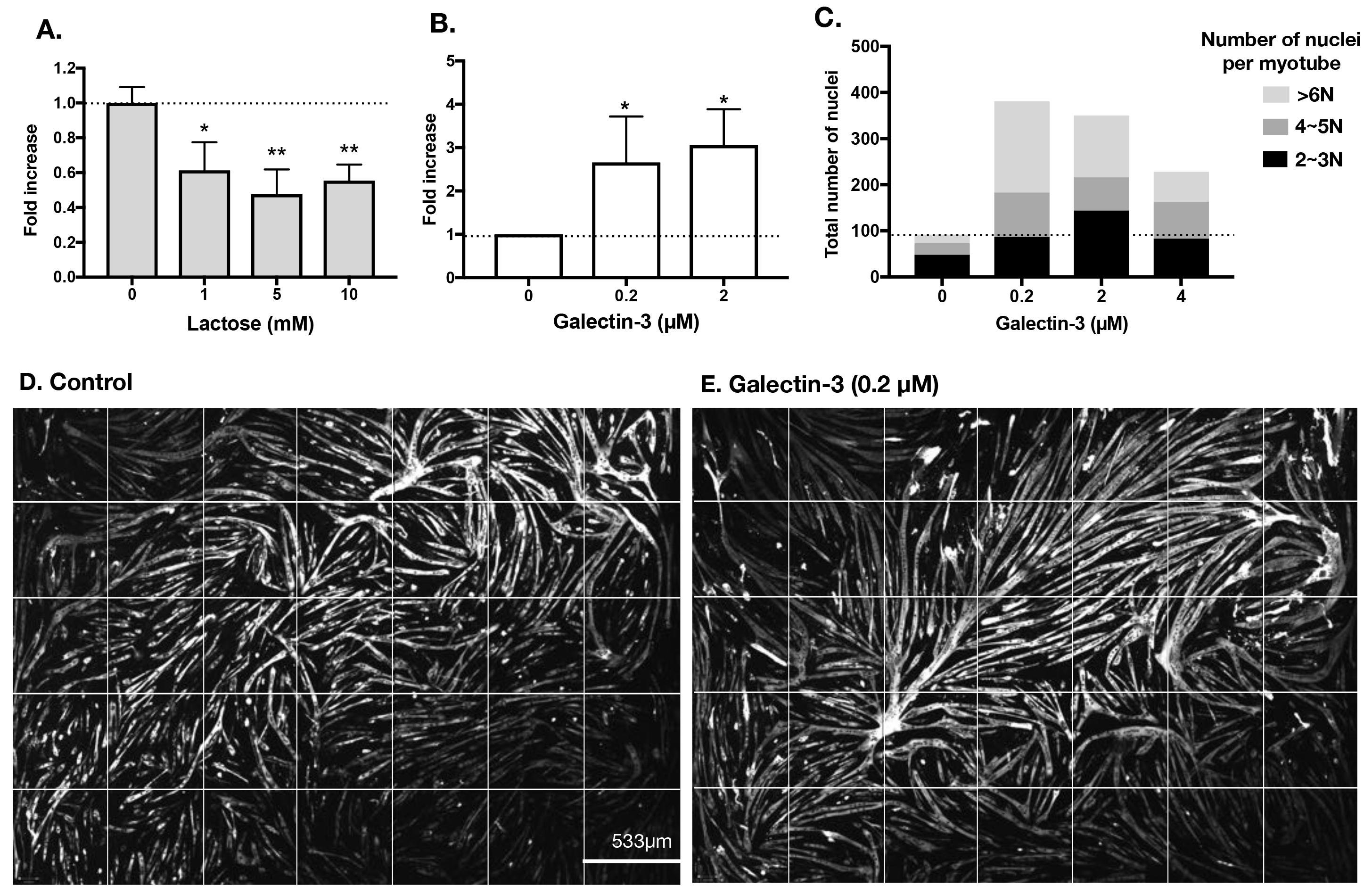
Galectin-3 promotes the myogenesis. **A**. Myogensis was induced in the presence or absence of different concentrations of lactose, an antagonist of galectin-3 for 72h. Cells were fixed and stained with anti-MHC antibody and DAPI. The total areas of MHC-positive multinucleated myotube were calculated by using in-house software. **B and C**. Myogenesis was induced in the presence or absence of galectin-3 for 62 h (B) and 72 h (C). The numbers of MHC-positive and multinuclear myotubes were counted. In C, total number of nuclei was counted and categorized into 3 groups based on the number of nuclei in each myotube. **D and E**. Stitched images (7 × 5 FOVs) of extended-focus images of myotubes (MHC-positive cells are in white) formed in the absence (D) and presence of galectin-3 (E) after differentiation for 72 h, are shown together with the size of one FOV. Significance compared to control (PBS-treated) condition * P < 0.05, ** P < 0.01

### Monosaccharide GlcNAc, which increases the biosynthesis of the glycan ligands of galectin-3, increases myogenesis

Since galectin-3 increased the efficiency of myogenesis, we next tested whether the myogenesis is also enhanced by increasing the expression of galectin-3 ligands. Most galectin-3 ligands are a specific type of asparagine-linked oligosaccharide on membrane glycoproteins (30). These oligosaccharides carry many *N*-acetyllactosamine residues (LacNAc, Galβ1-4GlcNAc) on their branches (Fig. 3A) and the affinity and avidity of galectin-3 for these oligosaccharides increases in proportion to the number of LacNAc units (31, 32). The number of LacNAc units strongly depends on the number of GlcNAc branches, especially β1,6 branches, the synthesis of which is mediated by the enzyme MGAT5 (α1,6-mannosyl glycoprotein 6-βGlcNAc transferase) (31, 32). Importantly, MGAT5 catalyzes the rate-limiting step in the synthesis of the oligosaccharide ligands of galectin-3. MGAT5 transfers GlcNAc from UDP-GlcNAc to the precursor oligosaccharides attached to proteins in the Golgi apparatus, but the reaction is suboptimal because the Michaelis constant (K_m_) of MGAT5 for UDP-GlcNAc (11 mM) is around 10-fold greater than the intra-Golgi UDP-GlcNAc concentration (32). Therefore, an increase in the UDP-GlcNAc concentration enhances the synthesis of galectin-3 ligands in a linear manner. The intracellular concentration of UDP-GlcNAc can be increased by increasing the extracellular concentration of GlcNAc (Fig. 3A) (31). Previous studies using immune and cancer cells demonstrated *in vitro* and *in vivo* that the treatment of cells with GlcNAc increases the expression of galectin-3 ligands (15). Therefore, we next examined whether increase in the expression of galectin-3 ligands by GlcNAc promotes the myogenesis. As little as 0.2 mM GlcNAc was sufficient to increase myogenesis (Fig. 3B). In contrast, a control saccharide, mannose (10mM), had no effect on the efficiency of myogenesis, while a weak antagonist of galectin-3, galactose, inhibited the myogenesis at 10mM but not 5mM (Fig. 3C and data not shown). Similar to galectin-3, GlcNAc increased both the length and thickness of the myotubes (Fig. 3D and E) and also promoted the formation of myotubes that contain many nuclei (Fig. 3F). An antagonist of galectin-3, lactose, inhibited GlcNAc-induced myogenesis (Fig. 3G).

**Fig. 3.**
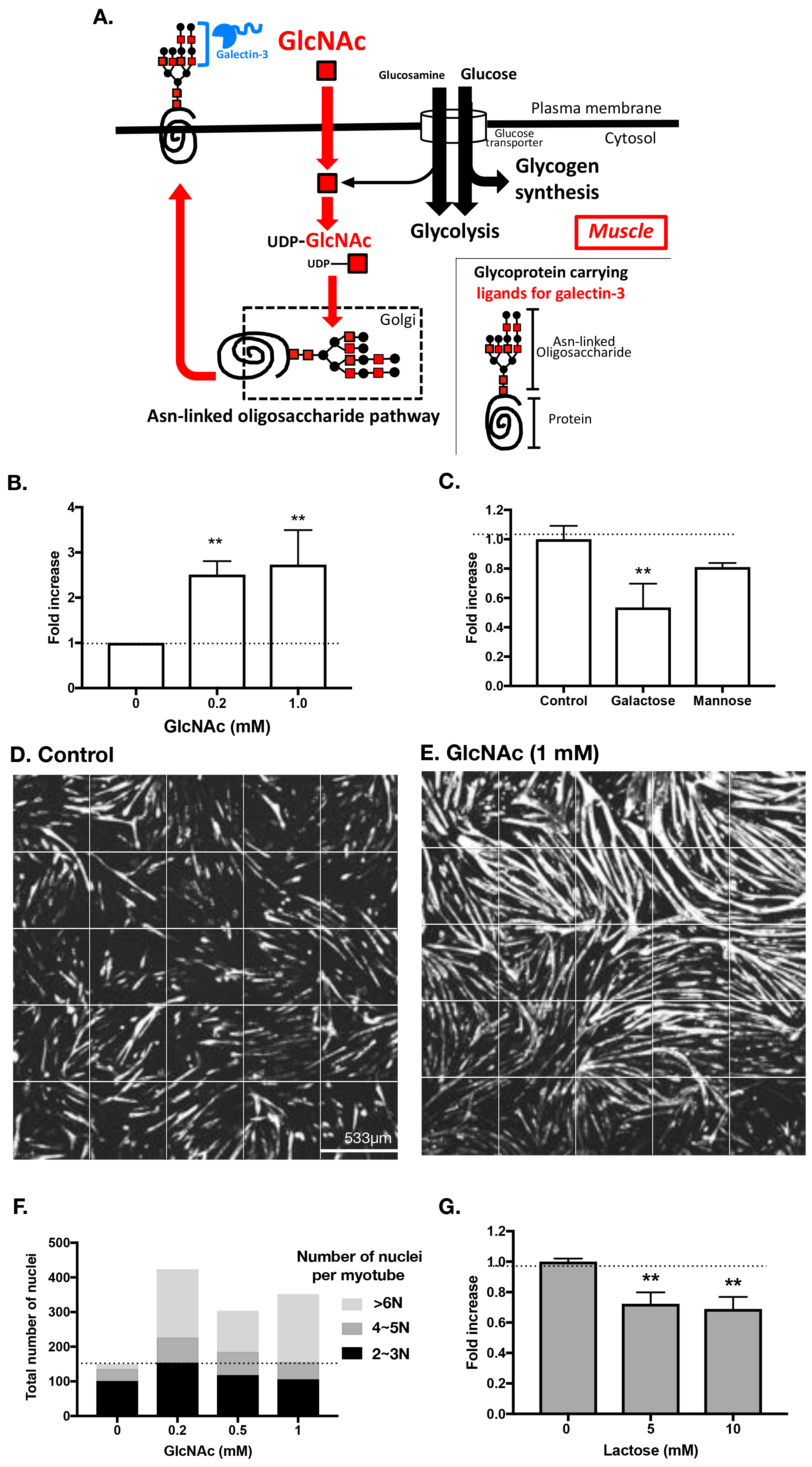
GlcNAc promotes myogenesis. **A**. Regulation of the synthesis of asparagine-linked glycans by GlcNAc in muscle cells. Glucosamine is taken up by a glucose transporter. Most glucosamine is used for glycolysis and glycogen synthesis and only a small percentage is transformed to UDP-GlcNAc. In contrast, extracellular GlcNAc is rapidly endocytosed to synthesize UDP-GlcNAc. UDP-GlcNAc is then incorporated into asparagine-linked glycans, which are attached to proteins in the Golgi apparatus. The glycoproteins are then transported to the cell surface. Galectin-3 binds to the glycoproteins. **B**. Myogenesis was induced for 62 h in the presence or absence of GlcNAc. Numbers of myotubes and nuclei in the myotubes were counted. **C**. Myogenesis was induced in the presence or absence of saccharides (galactose and mannose) at the concentration of 10 mM. The total areas of MHC-positive multinucleated myotubes were calculated. **D and E**. Stitched extended-focus images (5 × 5 FOVs) of myotubes (MHC-positive cells are white) formed in the absence (D) or presence of GlcNAc (E) after differentiation for 72 h. **F**. Myogenesis was induced for 72h and total number of nuclei was counted and categorized into 3 groups based on the number of nuclei in each myotube **G**. Myogenesis was induced in the presence of GlcNAc together with different concentrations of the galectin-3 antagonist, lactose. *P < 0.05, **P < 0.01.

### Lack of galectin-3 impairs the force production capacity of soleus muscles

To examine the role of galectin-3 in muscle physiology, we compared the maximum specific force in both wild-type mice and galectin-3-deficient mice. As shown in Fig. 4, the specific tetanic tension of the soleus muscles in the galectin-3-deficient mice was inferior to that in the wild-type mice, suggesting that the lack of galectin-3 impairs the capacity for muscle force production.

**Fig. 4.**
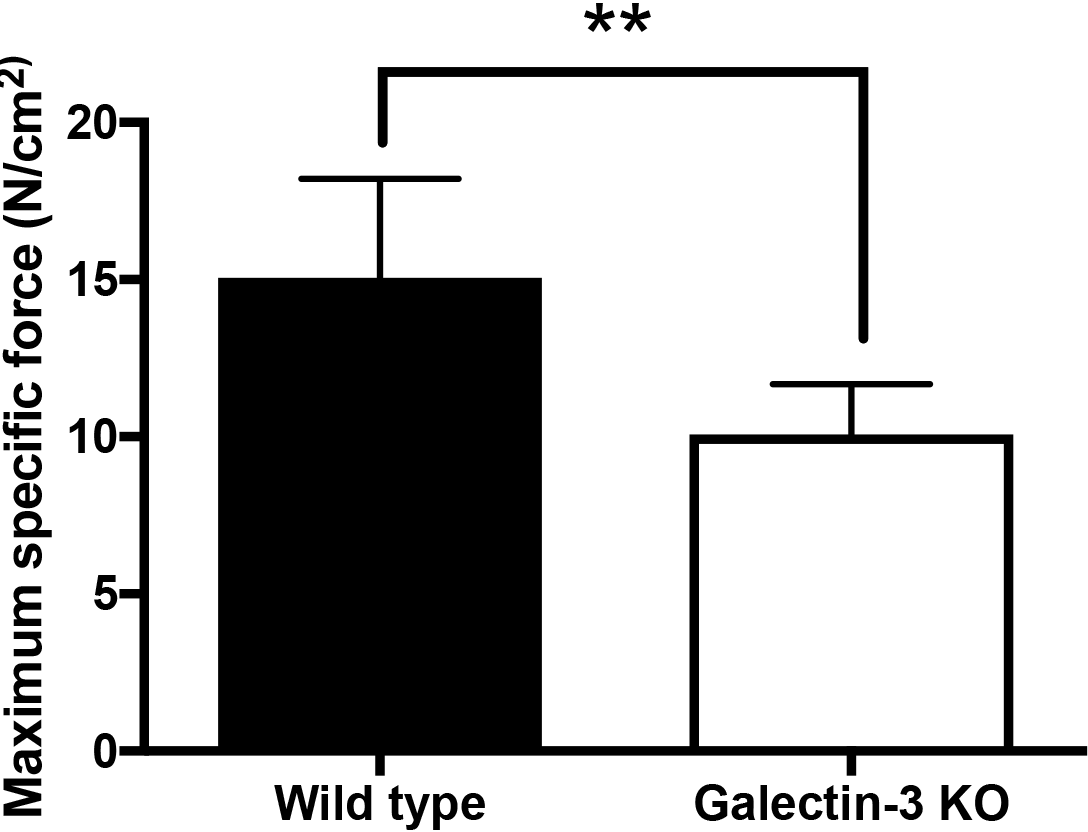
Lack of galectin-3 impairs the force production capacity of soleus muscles. Soleus muscles were isolated from galectin-3-knockout mice and the maximum specific force was measured. **P < 0.01.

### GlcNAc treatment increases muscle function in *mdx* mice

We next assessed the therapeutic potential of GlcNAc in DMD model mice. *Mdx* mice were intraperitoneally administered either PBS or GlcNAc daily, at a dose of 250 mg/kg bodyweight, for 10 days starting from 25 days after birth, i.e. the first and most important peak of muscle degeneration in this mouse model. Treatment with GlcNAc significantly reduced the damaged areas in the EDL muscles (Fig. 5A and B). The force-frequency curves and maximum specific force of the fast-twitch fibers of the isolated EDL and the slow-twitch fibers of the isolated soleus muscles were tested after treatment for 10 days (Fig. 5C-F). No significant difference was observed in body weight, muscle weights, peak tension, half relaxation time and twitch tension (Supplementary Table S1). At low frequencies, the force-frequency curves were similar between GlcNAc-treated and control (Fig. 5C and D). In contrast, at high frequencies, GlcNAc significantly increased force production of both EDL and soleus muscle relative to PBS-treated control (Fig. 5C and D). Maximum specific force generation of both EDL and soleus muscle was also significantly higher in mice treated with GlcNAc compared to that in the PBS-treated mice. Those results suggest that treatment with GlcNAc significantly improved the functions of dystrophic muscles.

**Fig. 5.**
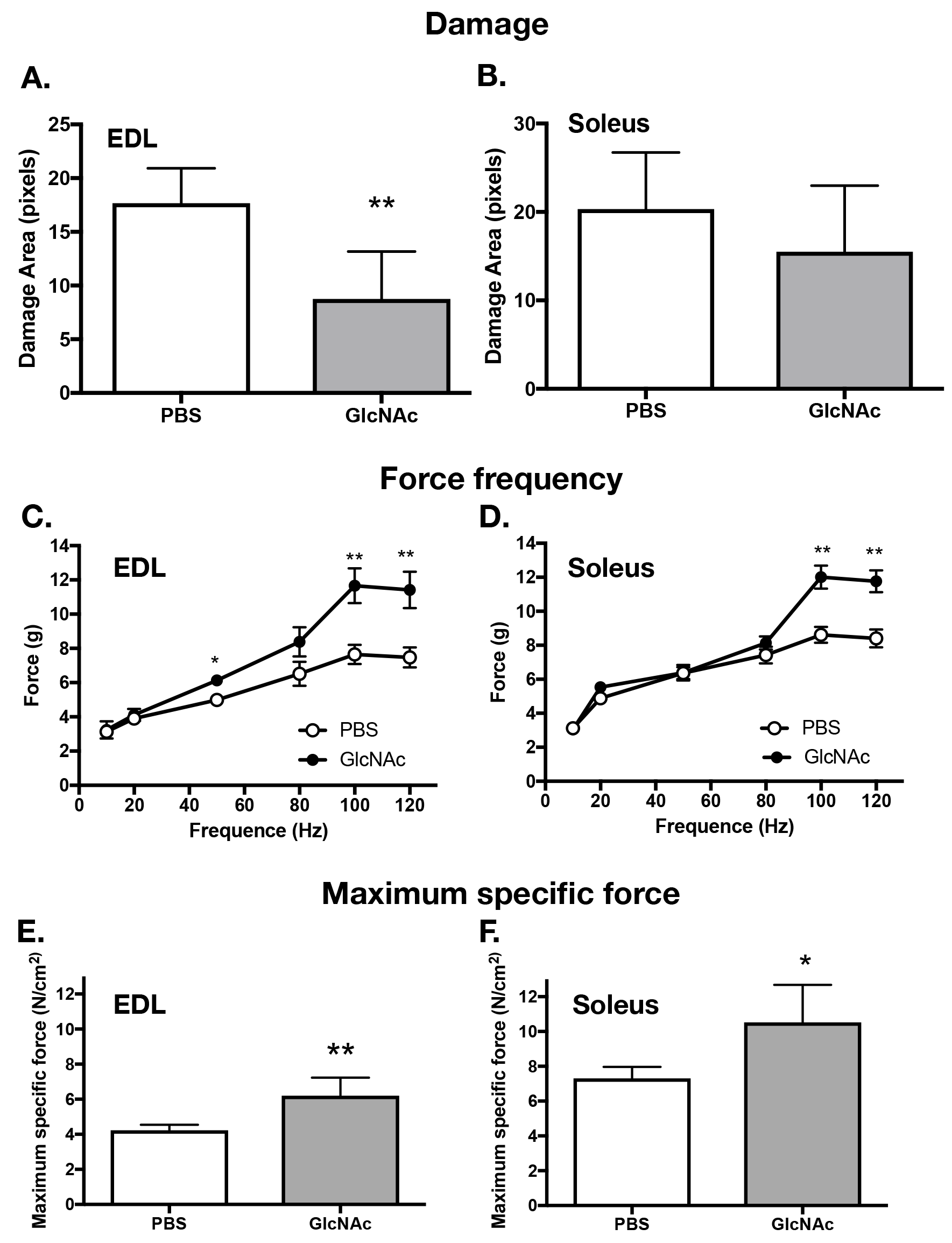
GlcNAc treatment improves the function of dystrophic muscles in *mdx* mice. Dmd^mdx^ mice were treated intraperitoneally with GlcNAc (25 mg/kg bodyweight/day) for 10 days, starting from 25 days after birth. A and B. Soleus and EDL muscles were collected on day 35 and stained with hematoxylin and eosin. The areas of muscle damage were estimated. C-F. Soleus and EDL muscles were collected on day 35 and their isometric contractile properties (C-D), and maximum specific force (E-F) were measured. *P < 0.05, **P < 0.01.

## Discussion

In this study, we have shown that galectin-3 or the monosaccharide GlcNAc, which increases the biosynthesis and expression of the glycan ligands of galectin-3, increases the efficiency of myogenesis. We have also presented evidence in a mouse model of DMD that GlcNAc has therapeutic potential in rescuing the functions of both slow-and fast-twitch dystrophic skeletal muscles during the initial peak of muscle degeneration. It has been reported that mice lacking the key enzyme MGAT5, which is the rate-limiting enzyme in the biosynthesis of galectin-3 ligands, show impairment of muscle repair and growth with aging (33). Our results demonstrate that the maximum specific force of the soleus muscles in galectin-3-deficient mice was inferior to that in wild-type mice, suggesting that the lack of galectin-3 impairs the capacity to generate force. Together, this evidence suggests that galectin-3 plays a critical role in muscle biology by interacting with its glycan ligands. These observations raise the possibility that a noninvasive therapy for DMD can be developed using GlcNAc.

The molecular mechanisms by which the interaction between galectin-3 and its glycan ligands promotes myogenesis and mitigates MD are still unknown. Once galectin-3 molecules bind to their ligand proteins, they cross-link those proteins. Previous studies in other tissues have suggested that galectin-3 mediates cell-cell interactions, such as the adhesion of neutrophils to endothelial cells and to laminin (14, 16, 34). Therefore, it is possible that galectin-3 facilitates the intimate interactions of differentiating myoblasts that promote myogenesis. Immediately before the fusion of those adherent myoblasts, phosphatidylserine (PS), a phospholipid that localizes exclusively in the inner cytoplasmic leaflet of membranes, is transiently exposed on the surface of the fusing membranes, and this transient exposure of PS is required for cell fusion (35). Interestingly, galectin-3 reportedly induces the transient exposure of PS on some cells, including lymphocytes and neutrophils without inducing cell apoptosis (36). Galectin-3 also binds to the insulin-like growth (IGF) receptor and positively modifies its signal transduction in microglia cells (11), and this IGF pathway also regulates myogenesis (18, 19). Therefore, it is possible that galectin-3 is involved in several differentiation steps in myogenesis. Dystrophic muscles are subject to repeated injury because the sarcolemmal membrane does not stably adhere to the basal lamina. Galectin-3 is one of the laminin-binding proteins, and this binding is also considered to enhance cell adhesion to the basal lamina. It is possible that the observed reduction in muscle injury in *mdx* mice treated with GlcNAc is also related to the stabilization of sarcolemmal membrane adhesion to laminin. Muscle injury induces the recruitment of proinflammatory macrophages. Once recruited, some of these switch phenotype to M2-type regulatory macrophages, which promote myogenesis (8, 37, 38). Interestingly, galectin-3 is involved in the maintenance of the M2 phenotype of macrophages (6). Therefore, it is also possible that both galectin-3 and a treatment that increases the biosynthesis of galectin-3 ligands positively affect the health of dystrophic muscles by several routes. The mechanisms by which GlcNAc mitigates MD warrant further investigation to facilitate the development of a novel therapy for the treatment of DMD.

GlcNAc is related to glucosamine, a compound that is believed to reduce arthritic pain, although its underlying molecular mechanism remains totally unknown. Importantly, the biological effect of GlcNAc is distinct from that of glucosamine (39). Glucosamine is entered to cells through glucose transporters and more than 95% of it is used for glycolysis, and in the case of muscles, also for glycogen synthesis (Fig. 3A). In contrast, GlcNAc is taken up by endocytosis and most is then rapidly converted to UDP-GlcNAc, which is incorporated into asparagine-linked glycans on membrane proteins (Fig. 3A). Previous studies using immune and cancer cells have demonstrated *in vitro* and *in vivo* that the treatment of cells with high concentrations of GlcNAc (10-20 mM) increases the expression of galectin-3 ligands (9, 40). Interestingly, we found that a as 0.2 mM GlcNAc was sufficient to promote myogenesis. This result suggests the possibility that the biosynthesis of glycan ligands can be efficiently modified by the administration of GlcNAc to muscle cells, where most glucose and glucosamine is consumed in glycolysis and/or glycogen synthesis rather than converted to UDP-GlcNAc for glycan biosynthesis. Because muscle tissue accounts for approximately 40-45% of the human body mass, the administration of the monosaccharide GlcNAc as a mitigating therapy for MD is an interesting therapeutic option. Importantly, GlcNAc does not exert any major adverse effects in humans, even at a dose of 25 mg/kg bodyweight/day for 6 weeks (41) or 6 g daily in children (42). The chronic administration of GlcNAc at 2.5 g/kg bodyweight/day to rats for 52 weeks induced no apparent adverse effects or histopathological changes in their tissues (43, 44).

In this study, we present the evidence that galectin-3 and GlcNAc, which increases the level of galectin-3 ligands, have interesting therapeutic potential to mitigate some symptoms associated with DMD. While further study is essential to understand the mechanism by which both galectin-3 and GlcNAc mitigate DM and promote myogenesis, the present study indicates that using GlcNAc as a supplement agent may present an interesting class of therapy for DMD, especially, the safety of this inexpensive monosaccharide is relatively established in humans.

## Author Contributions

S. Sato and J. Frenette designed research, analyzed data and wrote the paper. A. Rancourt, S. Dufresne, G. St-Pierre, H. Nakamura and S. Sato performed research. Y. Kikuchi and J. C. Lévesque analyzed data. M. S. Satoh developed software necessary to analyze data.

## Acknowledgments

We would like to acknowledge the Bioimaging Platform at the Centre de Recherche CHU de Quebec. The monoclonal antibody directed against MHC was obtained from the Developmental Studies Hybridoma Bank, created by the NICHD of the National Institutes of Health and maintained at the Department of Biology, University of Iowa, Iowa City, IA 52242. We also thank the Consortium for Functional Glycomics for galectin-3KO mice.

